# Survival of mineral-bound peptides into the Miocene

**DOI:** 10.1101/2022.08.19.502663

**Authors:** Beatrice Demarchi, Meaghan Mackie, Ziheng Li, Tao Deng, Matthew J. Collins, Julia Clarke

## Abstract

The oldest authenticated peptide sequences to date were reported in 2016 from 3.8 Ma old ostrich eggshell (OES) from the site of Laetoli, Tanzania (Demarchi et al., 2016). Here we demonstrate survival of the same sequences in 6.5-9 Ma OES recovered from a palaeosteppe setting in northwestern China. The eggshell is thicker than those observed in extant species and consistent with the Liushu *Struthio* sp. ootaxon. These findings push the preservation of ancient proteins back to the Miocene and highlight their potential for paleontology, paleoecology and evolutionary biology.

## Introduction

The oldest authenticated peptide sequences to date were reported in 2016 from 3.8 Ma old ostrich eggshell (OES) from the site of Laetoli, Tanzania (Demarchi et al., 2016), with a thermal age of the peptides estimated to be equivalent to ~16 Ma at a constant 10°C. This finding had great scientific impact since it integrated computational chemistry (molecular dynamics simulations) as well as experimental data to propose a mechanism of preservation, concluding that mineral binding ensures survival of protein sequences. Importantly, this study demonstrated that peptide-bound amino acids could survive into deep time even in hot environments. This has fuelled the analysis of ancient proteins from other mineral matrices, namely tooth enamel (Cappellini et al., 2019; Welker et al., 2020, 2019)in order to reconstruct phylogenies.

The recovery of peptides has so far been limited to biomineral samples of Plio-Pleistocene age: our attempts at retrieving intact peptides from Cretaceous eggshell were unsuccessful, despite the fact that a genuine intracrystalline fraction of amino acids was preserved in the same sample (Saitta et al., 2020). Here we show that the same peptide sequences that exhibit strong binding to the calcite surface - and that were recovered from the Laetoli OES - also persist into the late Miocene, in a > 6.5-Ma-old eggshell sample from the Linxia Basin, northeastern Tibetan Plateau, China, Liushu Formation.

## Results and discussion

The ostrich eggshell (IVPP V26107) was recovered from an area between Xinji and Songming towns, Hezheng County, close to the border with Guanghe County, from mudstone facies of the Liushu Formation (Figure 1).

**Figure 1:**
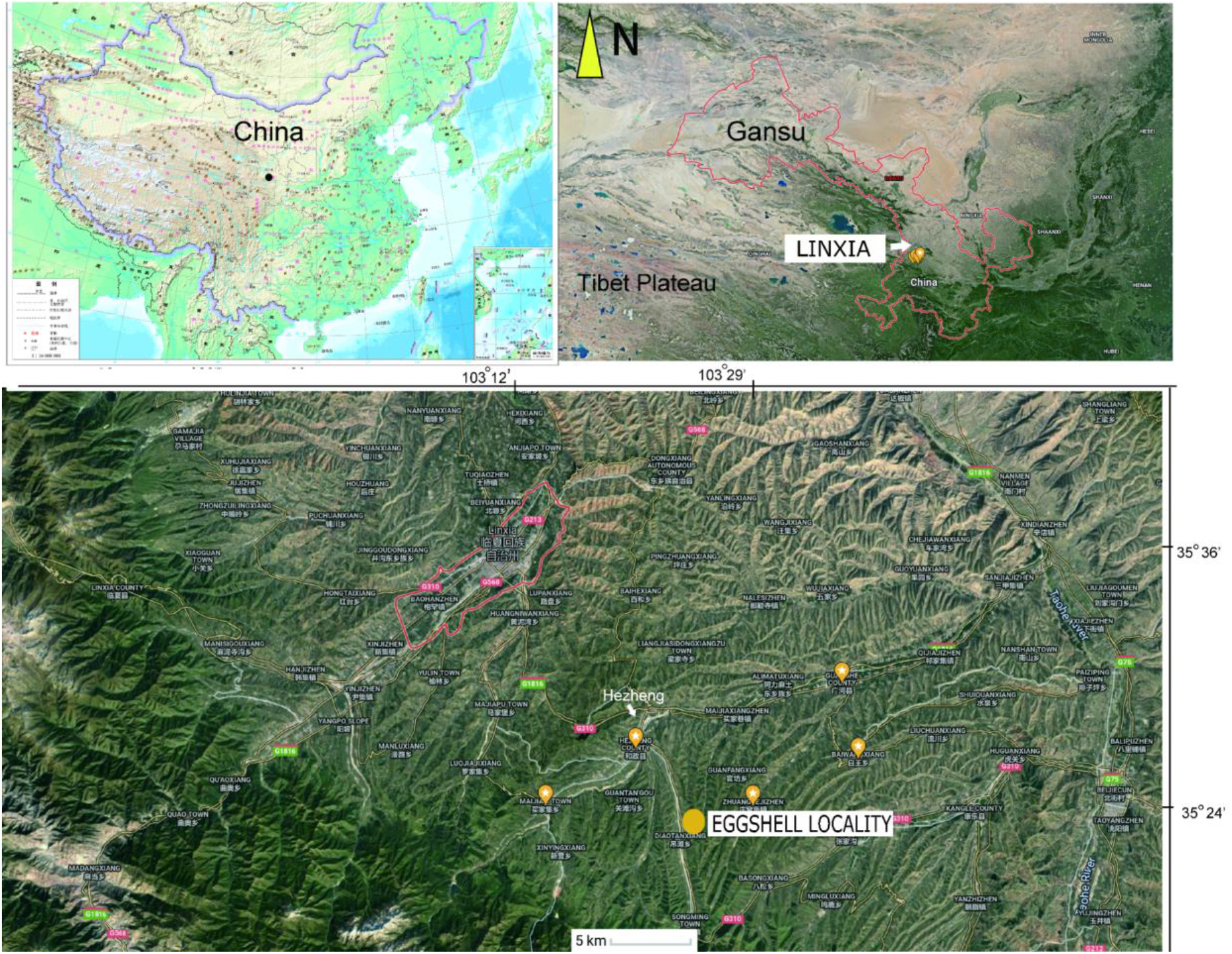
Map showing the location of the fossil eggshell site.

Both eggshell and skeletal remains are previously known from the Late Miocene Liushu Formation, Linxia Basin, of Gansu Province (Hou et al., 2005; Li et al., 2021; Wang, 2008). Age control on the highly fossiliferous units of the Lishu Formation is good (Deng et al., 2019, 2013; Zhang et al., 2019), with detailed correlations across China as well as with Neogene deposits globally. These units have been assessed via new magnetostratigraphic data; most of the *Hipparion* fauna is estimated to be 6.4-6.5 Ma, with a lower Liushu transition to a *Platybelodon* fauna yielding age estimates of 11.1-12.5 Ma (Zhang et al., 2019). These dates agree with the prior estimates from biostratigraphic and magnetostratigraphic data (Deng et al., 2019, 2013). The recovery site is closer to the southern part of the basin near the Heilinding section of Zhang et al. (2019), which had an estimated minimum age of 6.4-6.5 Ma (Chron 3An). In contrast, the candidate stratotype section, Guoniguo, that exposes lower parts of the Liushu, is north of the recovery area (Deng et al., 2019; Zhang et al., 2019).

Ostrich (*Struthio*) remains have a long history of recovery from the late Miocene of northwest China (Buffetaut and Angst, 2021; Hou et al., 2005; Li et al., 2021; Lowe, 1931; Mikhailov and Zelenkov,2020). The first reported ostrich fossils from China were from units of the *“Hipparion* Clay” (Red Clay) dated to between 6.54-7.18 Ma (Lowe, 1931; Zhu et al., 2008). Mikhailov and Zelenkov (2020) concluded that the Wang (2008) Liushu taxon is referable to *Struthio* and not presently supported as a distinct but related genus; it is proposed to be from a taxon larger than extant *Struthio* species with a thickness of ~2.4mm consistent with the eggshell sampled here (Figure 2C).

**Figure 2:**
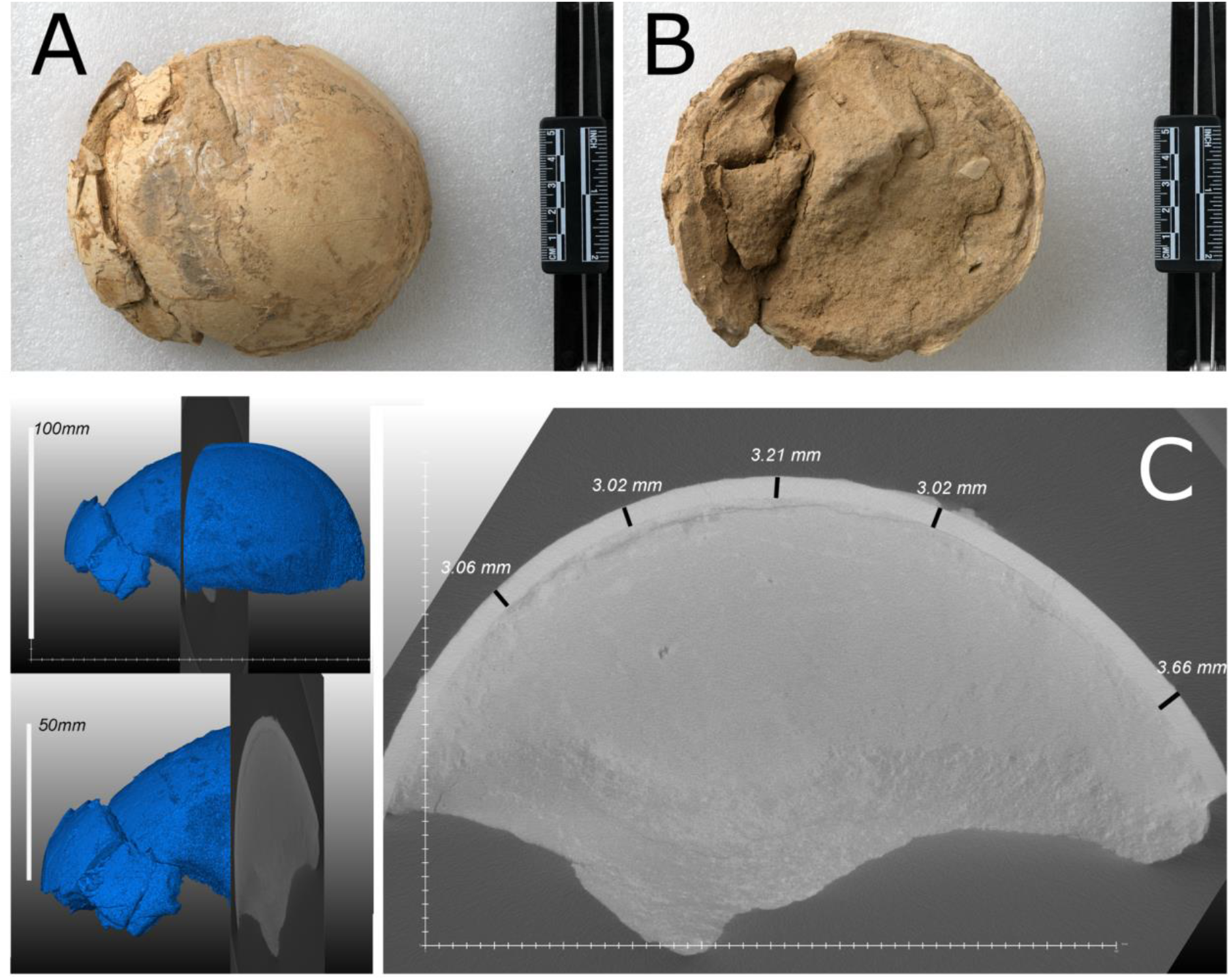
Eggshell specimen (IVPP V26107): photographs (A, B) and CT scan (C)

Thermal age calculations for the OES sample yielded a crude estimate of 6.8 Ma at a constant 10°C, thus thermal age and numerical age are broadly equivalent. Given the extreme antiquity of the samples, protein extraction was performed in an ultra-clean facility at the University of Copenhagen in order to minimise any chance of contamination, following Hendy et al. (2018). Furthermore, OES powders were bleached extensively to isolate the intracrystalline fraction only (as described by Demarchi et al. (2016)), samples were analysed by LC-MS/MS using a new LC column, and six blanks (analytical and procedural) were included in the run, flanking the sample. Data analysis was equally cautious: raw tandem mass spectrometry data were used to reconstruct potential peptide sequences starting from raw product ion spectra: 25 sequences were found with an ALC (Average Local Confidence) score ≥ 80%. This is the most stringent threshold that can be applied, and it signifies that all potential peptide sequences with lower confidence are discarded. Of these sequences, 14 contained the typical Asp-rich motif “DDDD” (Figure 3), which had been shown in our previous paper to be the mineral-binding peptide belonging to the sequence of struthiocalcin-1 (SCA-1).

**Figure 3:**
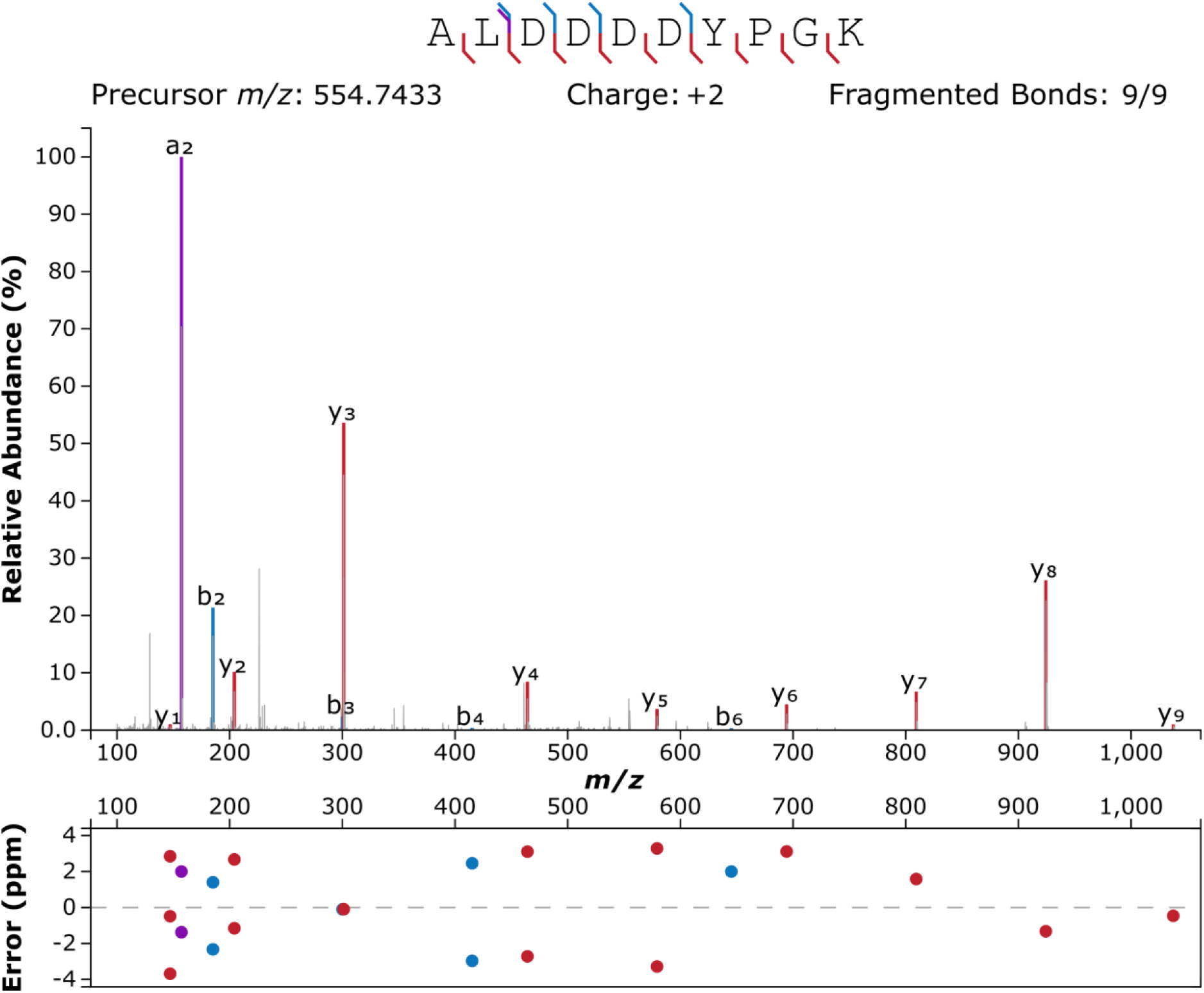
Annotated product ion spectrum of peptide ALDDDDYPGK. Figure created using www.interactivepeptidespectralannotator (Brademan et al., 2019).

The *de novo* peptides were searched against the Uniprot/Swissprot database (containing 565,254 manually annotated and reviewed protein sequences). A database of common laboratory contaminants (cRAP) was included in the search. The results were filtered according to the parameters detailed in the Methods section. The only protein identified was struthiocalcin-1. Figure 4 reports the coverage of SCA-1, including number of peptide-spectrum matches (PSM) and de novo tags (individual annotated tandem mass spectra are shown as Supplementary Figures).

**Figure 4:**
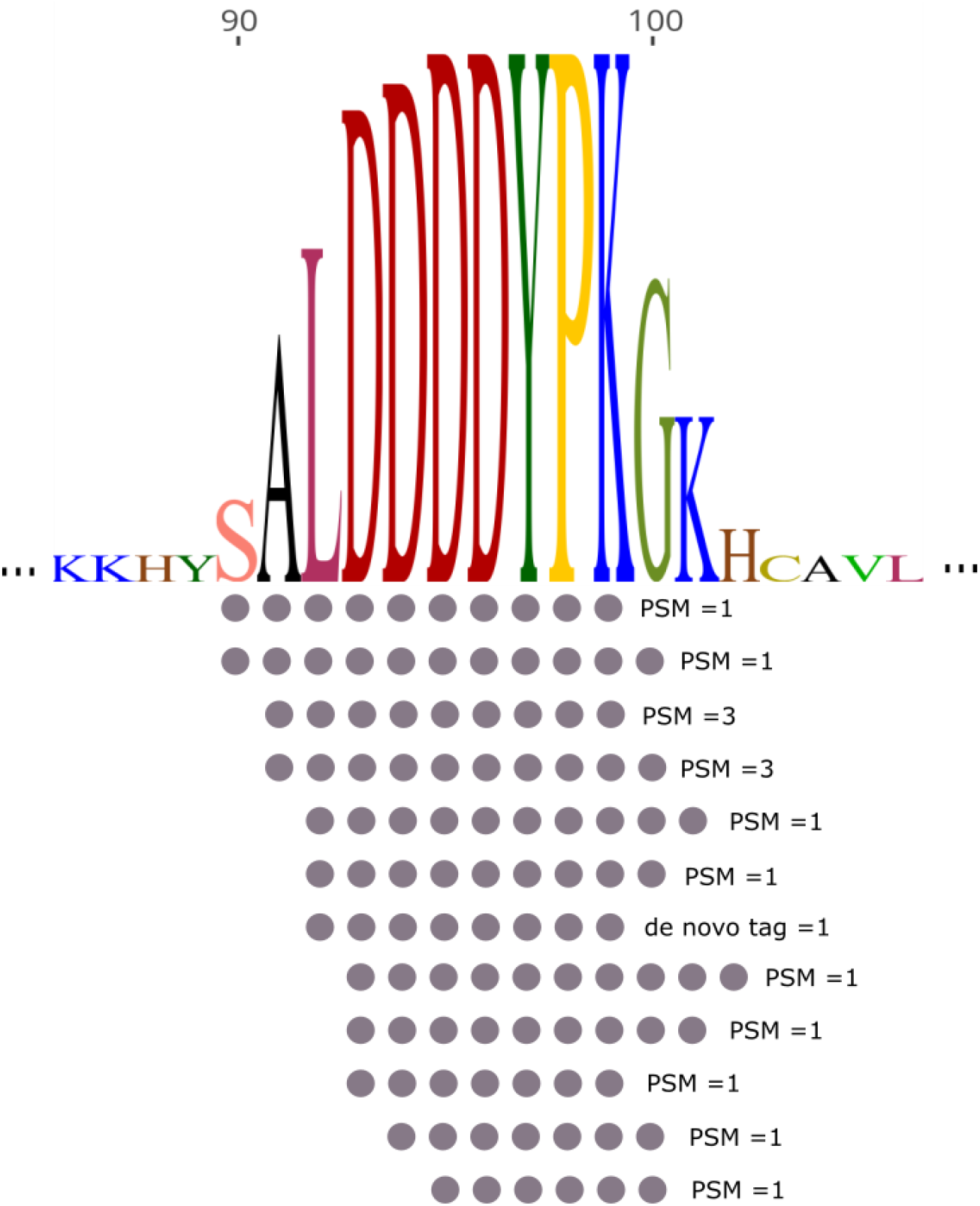
Coverage of struthiocalcin-1 (SCA-1) in Miocene OES specimen IVPP V26107.

## Conclusions

We present the first evidence for peptide survival into the Miocene, confirming our previous (Pliocene) data and providing further support to the mechanism of preservation based on the binding of Asx-rich peptides to calcite surfaces (Demarchi et al., 2016). The sequence recovered from the Chinese specimen is identical to the peptides found in the 3.8 Ma Laetoli OES, thus also supporting the attribution of the Liushu ootaxon to genus *Struthio.* While the four Asp residues are conserved across several avian taxa, in some species, including other ratites, Asp can be substituted by Glu and some of the flanking residues are also variable (Demarchi et al., 2022, fig. S1); therefore, variability within this sequence could be informative for evolutionary relationships between extinct and extant taxa.

## Materials and Methods

### Sample preparation and analysis

A subsample of OES specimen IVPP V26107 was prepared in the ultra-clean facility at the University of Copenhagen, following the protocol of Demarchi et al. (2022, 2016) and omitting the digestion step. In brief, the fragment was powdered, bleached for 72 hours (NaOCl, 15% w/v) and demineralised in just enough cold weak hydrochloric acid (0.6 M HCl). The extracts were exchanged in ammonium bicarbonate buffer (pH = 7.8) using 3kDa MWCO ultrafilters and the peptides purified and concentrated using C18 Stage Tips (Cappellini et al., 2019; Rappsilber et al., 2007).

Prepared StageTips of the procedural blank and sample were eluted with 30 μL of 40% acetonitrile (ACN) 0.1% formic acid into a 96-well plate prior to LC-MS/MS analysis. To remove the ACN, the plate was vacuum centrifuged until approximately 5 μL remained. Samples were then resuspended with 6 μL of 0.1% trifluoroacetic acid (TFA) 5% ACN. Based on protein concentration results at 205 nm (NanoDrop, Thermo Scientific), 5 μL of each sample and procedural blank was then separated over a 77 min gradient by an EASY-nLC 1200 (Proxeon, Odense, Denmark) attached to a Q-Exactive HF-X mass spectrometer (Thermo Scientific, Germany) using a 15 cm column. The column (75 μm inner diameter) was made in-house, laser pulled, and packed with 1.9 μm C18 beads (Dr. Maisch, Germany). Parameters were the same as those already published for historical samples (Mackie et al., 2018). In short, MS1: 120k resolution, maximum injection time (IT) 25 ms, scan target 3E6. MS2: 60k resolution, top 10 mode, maximum IT 118 ms, minimum scan target 3E3, normalized collision energy of 28, dynamic exclusion 20 s, and isolation window of 1.2 *m/z.* Wash-blanks consisting of 0.1% TFA 5% ACN were also run in order to hinder cross-contamination. The LC-MS/MS run included, in this order: two wash blanks, one procedural blank, one wash blank, the OES sample, and two wash blanks. Data are available via ProteomeXchange with identifier PXD035872.

### Data analysis

Bioinformatic analysis was carried out using PEAKS Studio 8.5 (Bioinformatics Solutions Inc (Zhanget al., 2012)). The Uniprot_swissprot database (downloaded 11/08/2021) was used for carrying out the searches and common contaminants were included (cRAP; common Repository of Adventitious Proteins: http://www.thegpm.org/crap/). No enzyme was specified for the digestion and the tolerance was set to 10 ppm on the precursor and 0.5 Da on the fragments. The thresholds for peptide and protein identification were set as follows: peptide score −10lgP ≥ 30, protein score −10lgP ≥ 20, *de novo* sequences scores (ALC%) ≥ 80.

### Thermal age estimates

We assumed an age for the eggshell to be Late Miocene, 6.4-6.5 Ma, in other words at the very youngest of the estimated age range of the Lishu Formation. The formation lies in the Linxia Basin of Gansu Province, where the Mean Annual Temperature (MAT) is ~ 11°C (Liu et al., 2016). A Late Miocene age would place the birds in a period in which, following uplift of the Tibetan Plateau during a phase of climate cooling (Holbourn et al., 2018), there was an increase in the intensification of the East Asian monsoon associated with a northwestern expansion of the humid belt (Musser et al., 2019). Local range of lithological palaeobotanical, fauna and isotopic data discussed in Deng (2006) suggest dryer conditions and a shift from broad-leaved woodland to a more seasonal open grassland environment.

A compilation of proxies suggests a drop in MAT ~6 °C over High-Mountain Asia during the Last Glacial Maximum (Yan et al., 2018), while PMIP3 predicts a decrease of ~5 °C at the LGM, similar to the similar to drop in Global surface air temperature used by Holden et al. (2019) in their simulation of Pliocene–Pleistocene climate (giving an estimated MAT from Late Miocene of 14.8°C). Assuming a present day MAT of ~ 11°C and a temperature at LGM of 6°C, and using the temperature data from Holden et al. (2019), extending the end Pliocene to 6.4 Ma as the date of the eggshell, we can provide a very crude estimate of thermal age (i.e. age at a constant 10 °C) of 6.8 Ma. This effectively means that the additional warming in the Miocene is partly offset by Pleistocene cooling. Thus whilst this sample is almost twice as old as the sample from Africa (Demarchi et al., 2016), this estimate suggests it is little over half the thermal age of eggshell from Laetoli, Tanzania. For example, fossils on present day Ellesmere Island, Nunavut (Wang et al., 2017) using the same model would have a thermal age 50 times younger, suggesting the possibility of much greater antiquity of proteins in high latitude sites.

## Acknowledgements

B.D.’s work was supported by a “Young Researchers - Rita Levi Montalcini” grant from the Italian Ministry of University and Research. M.M. and M.J.C. were supported by the Danish National Research Foundation Award PROTEIOS (DNRF128). J.C. acknowledges the Alexander von Humboldt Foundation and the Jackson School of Geosciences and Z.L. and D.T. the Chinese National Science Foundation. We thank Prof. Jesper Velgaard Olsen at the Novo Nordisk Center for Protein Research for providing access and resources, which were also funded in part by a donation from the Novo Nordisk Foundation (Grant No. NNF14CC0001).

## Supplementary Images

**Figure S1.**
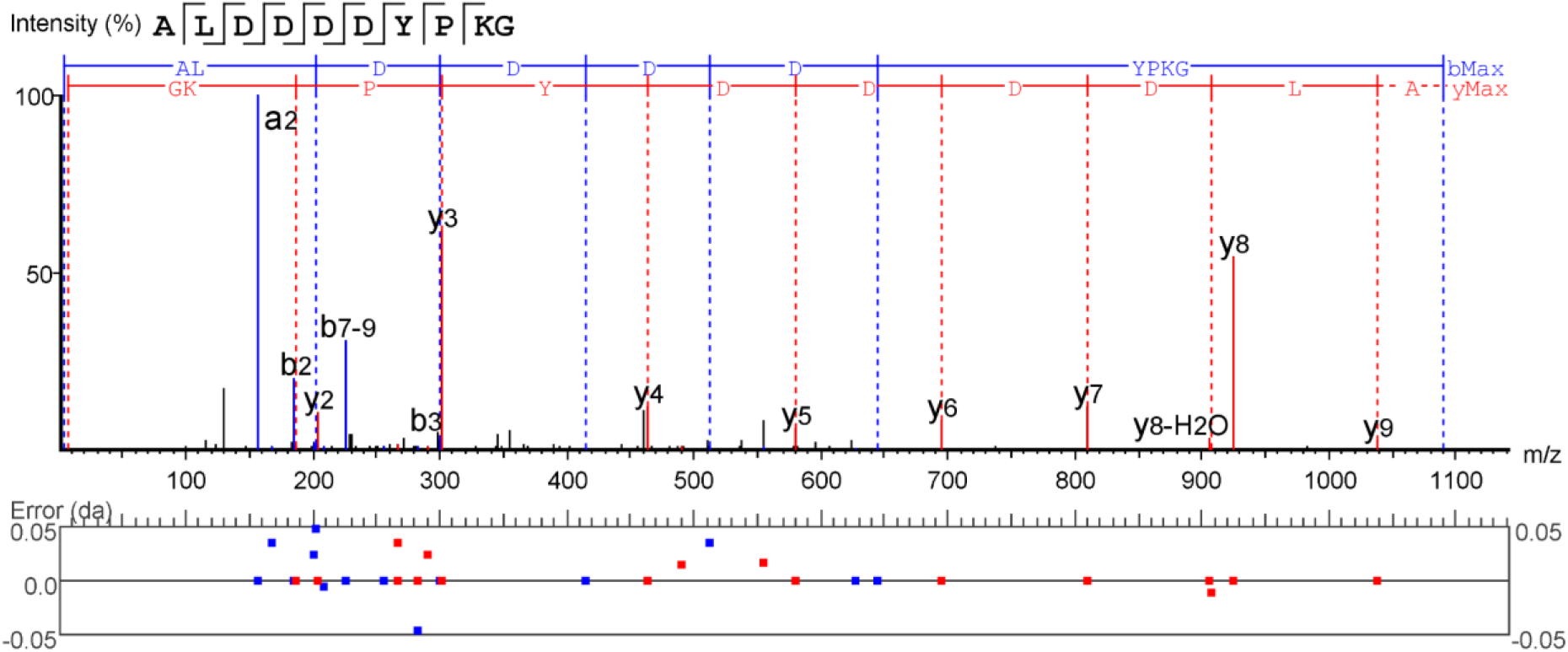
Annotated product ion spectrum, *m/z* = 554.7487.

**Figure S2.**
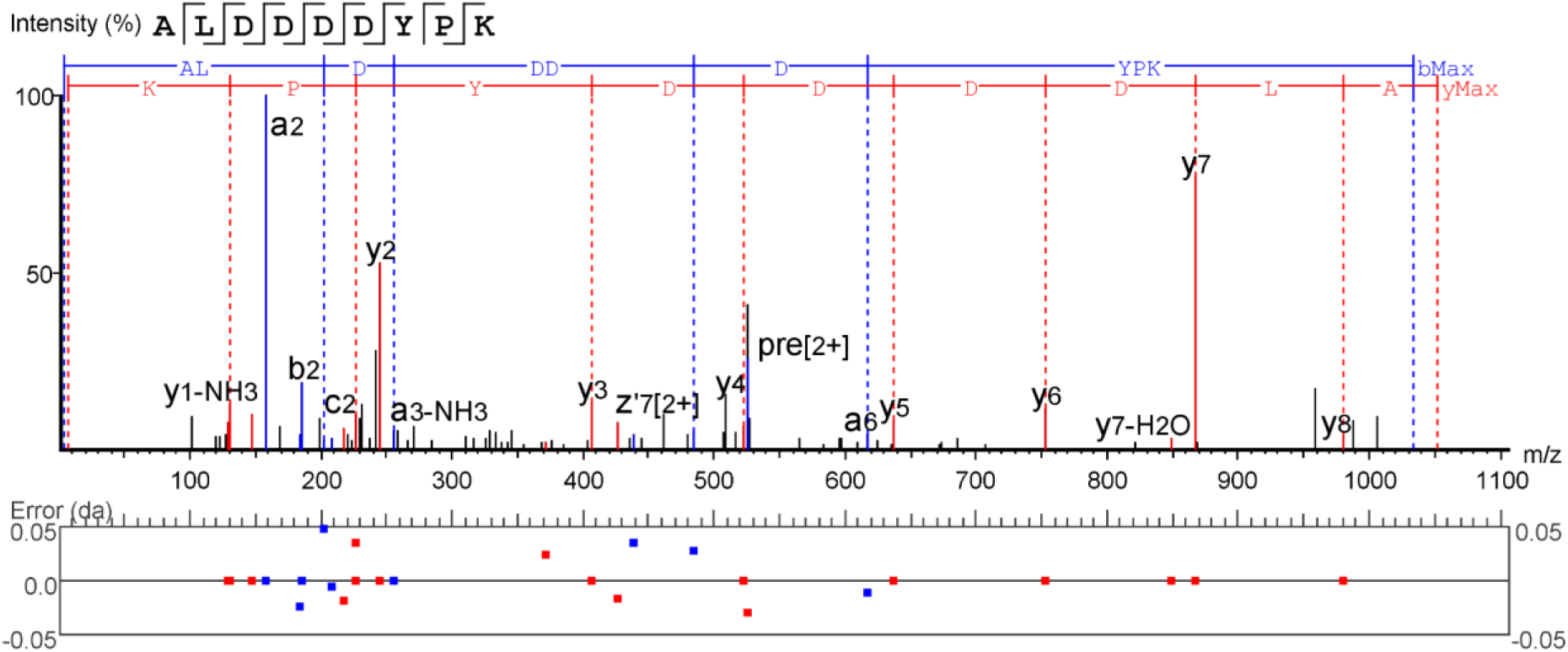
Annotated product ion spectrum, *m/z* = 526.2355

**Figure S3.**
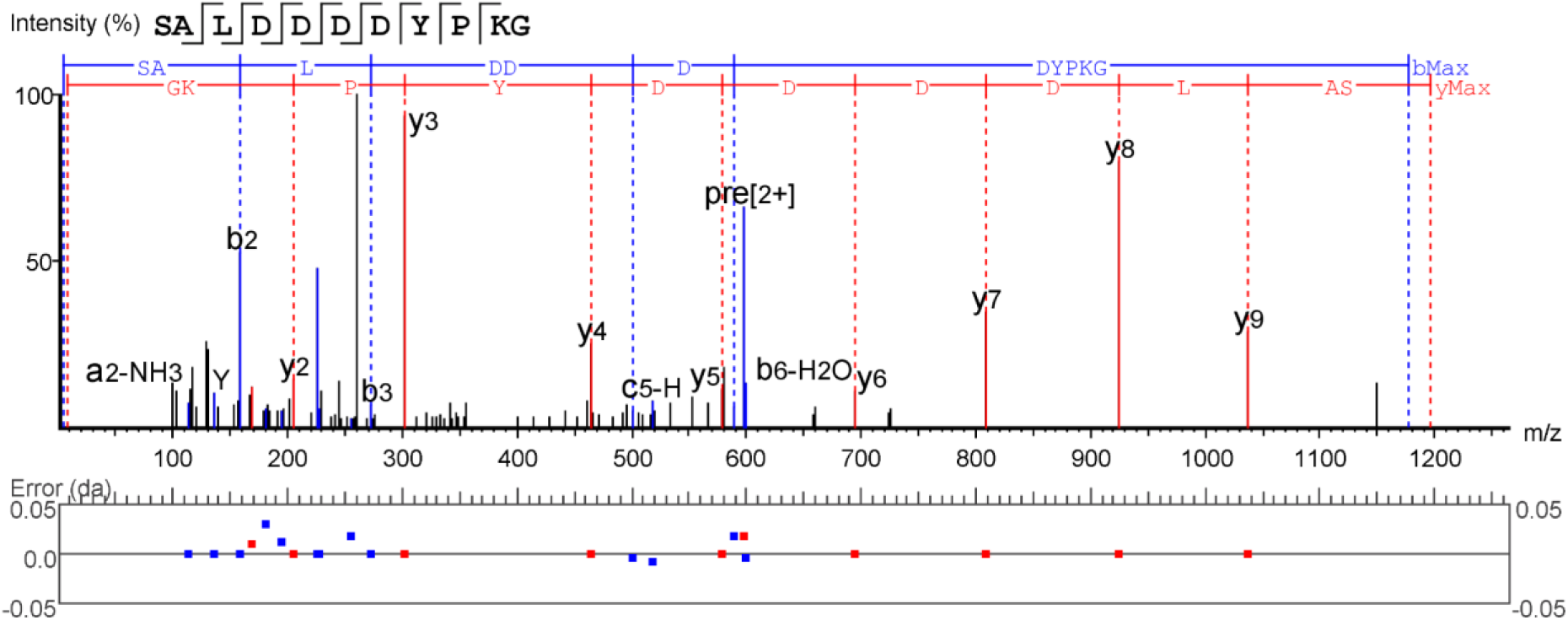
Annotated product ion spectrum, *m/z* = 598.2663

**Figure S4.**
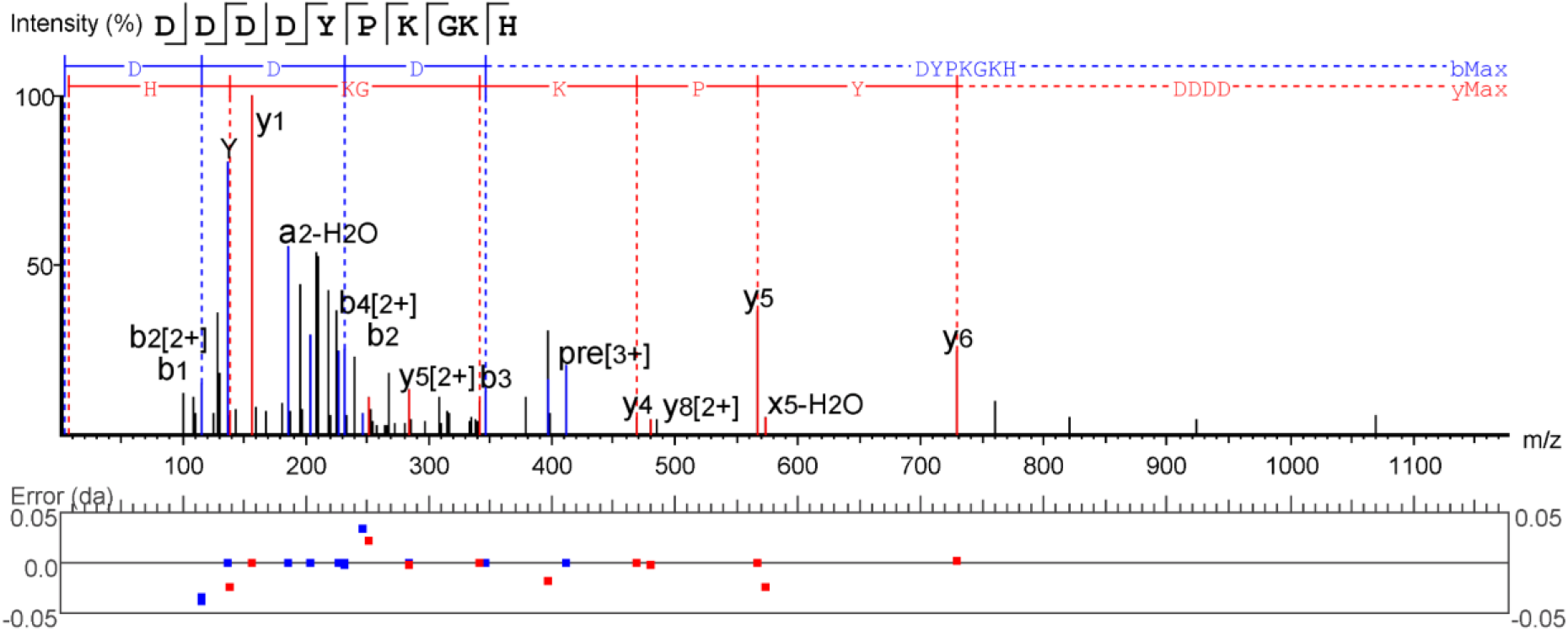
Annotated product ion spectrum, *m/z* = 397.1782

**Figure S5.**
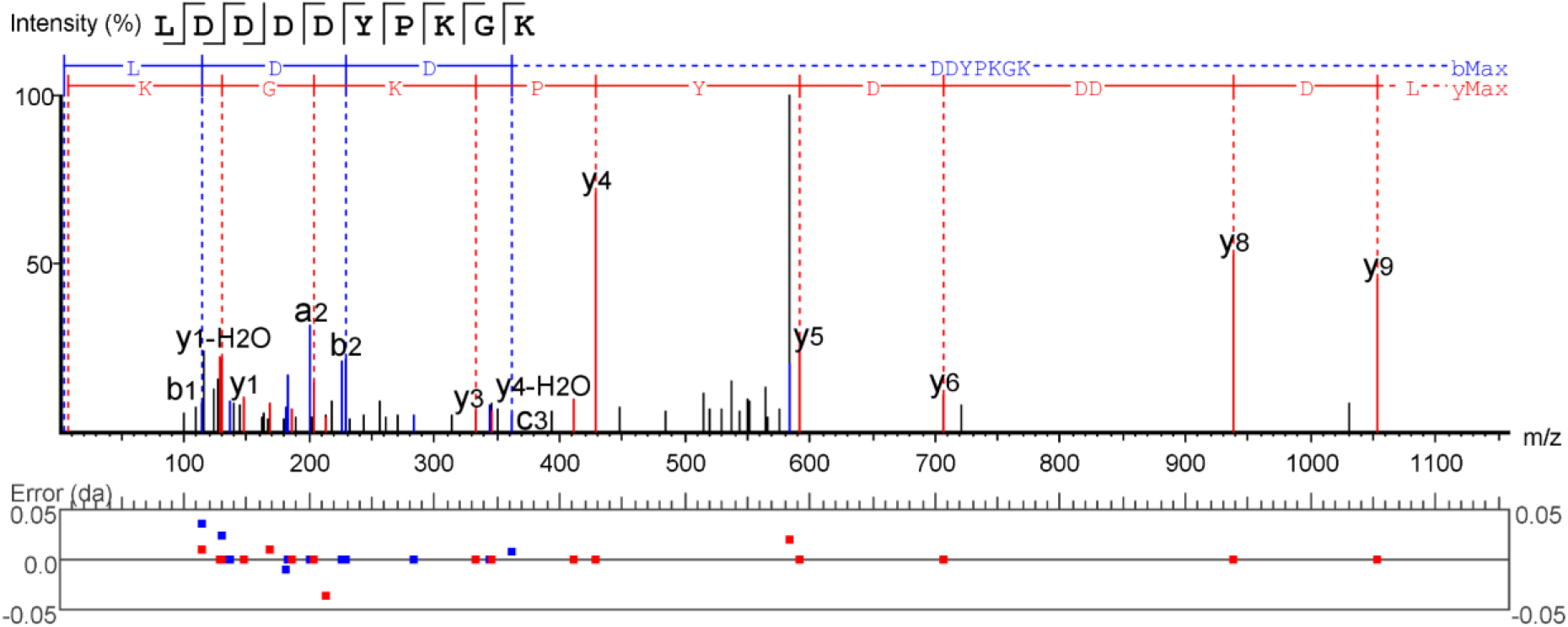
Annotated product ion spectrum, *m/z* = 583.2767

**Figure S6.**
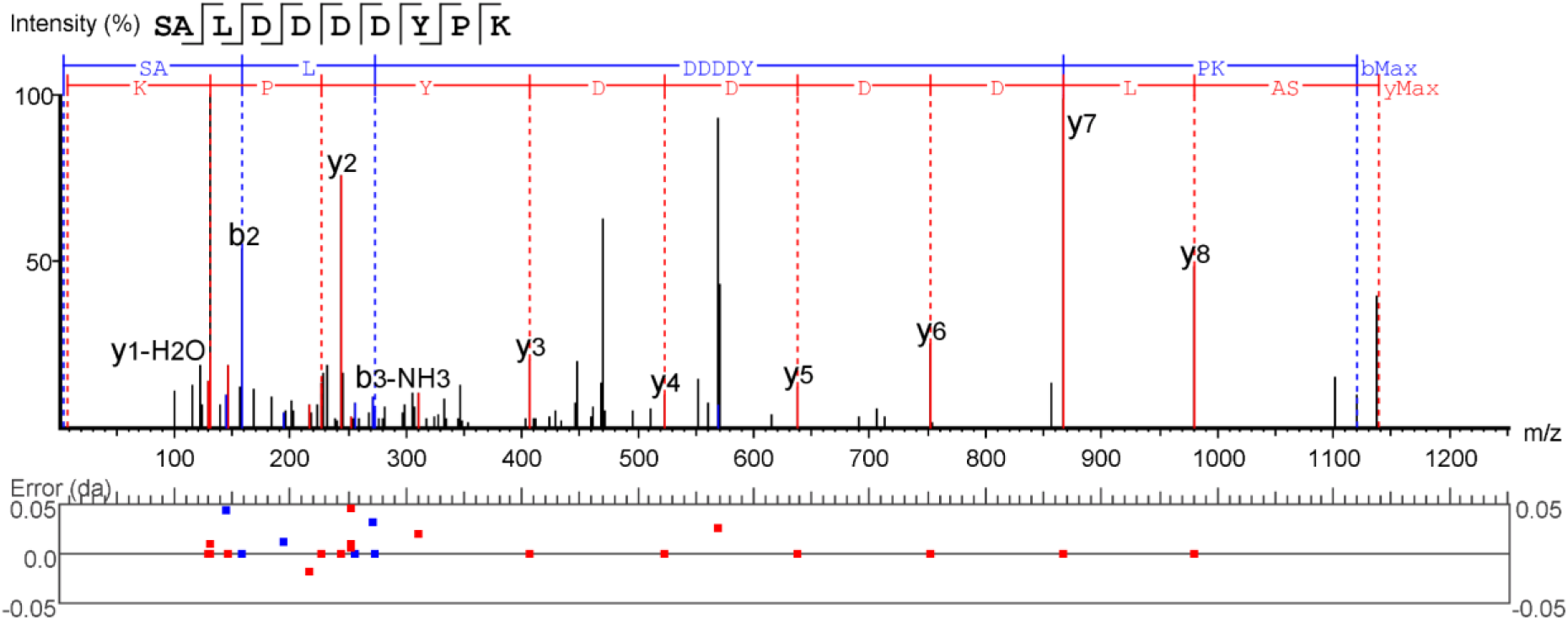
Annotated product ion spectrum, *m/z* = 569.7542

**Figure S7.**
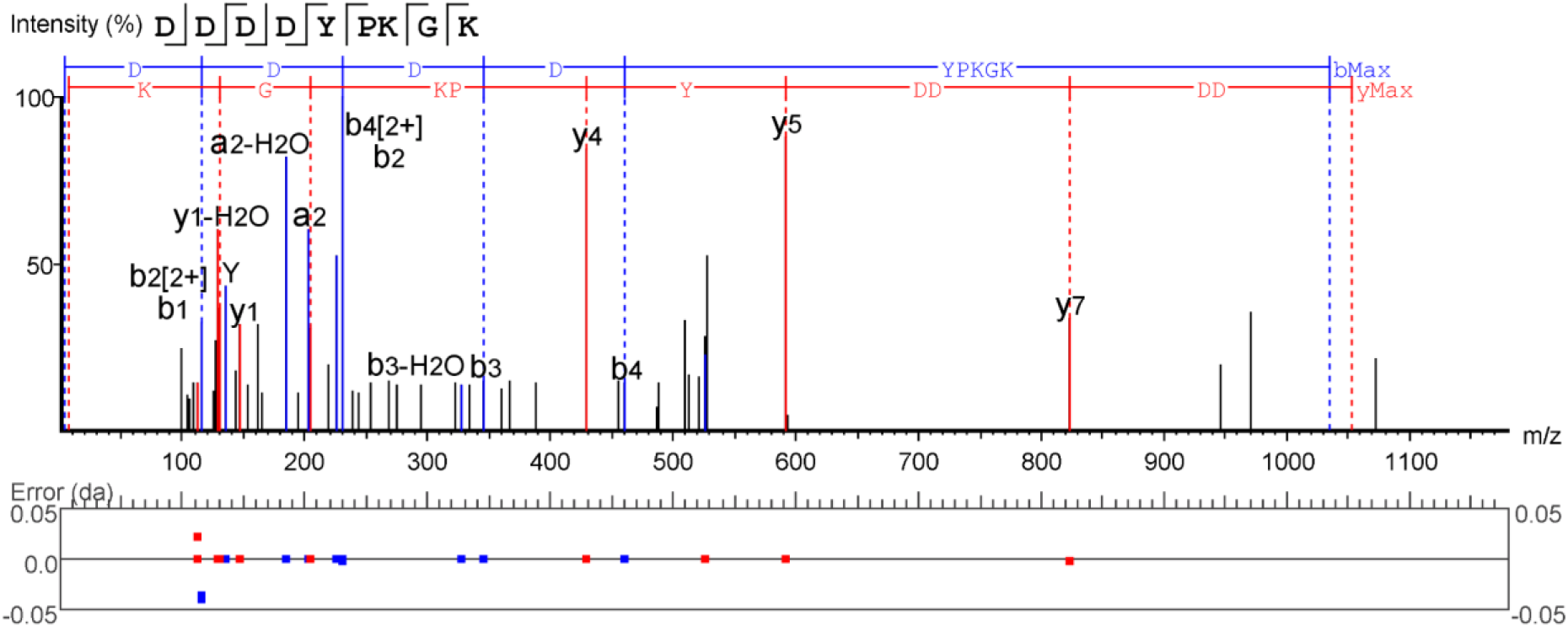
Annotated product ion spectrum, *m/z* = 526.7343

**Figure S8.**
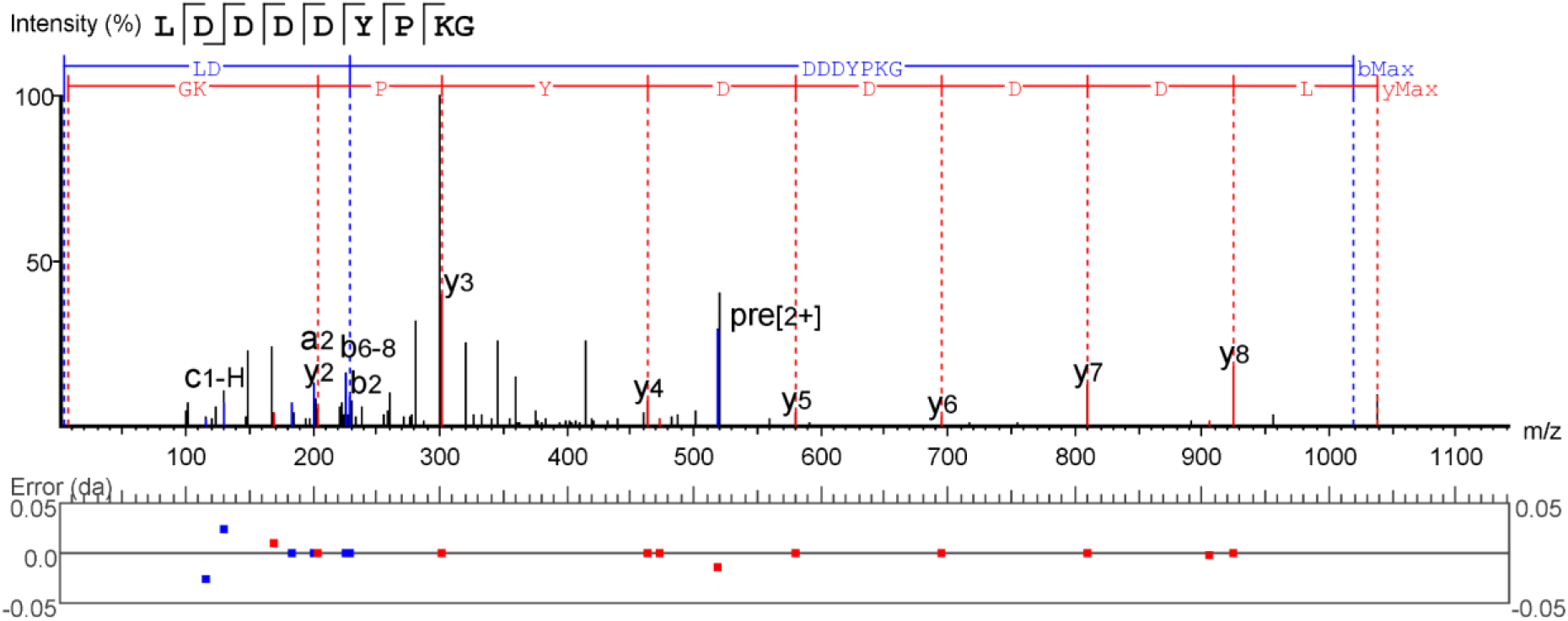
Annotated product ion spectrum, *m/z* = 519.2303

**Figure S9.**
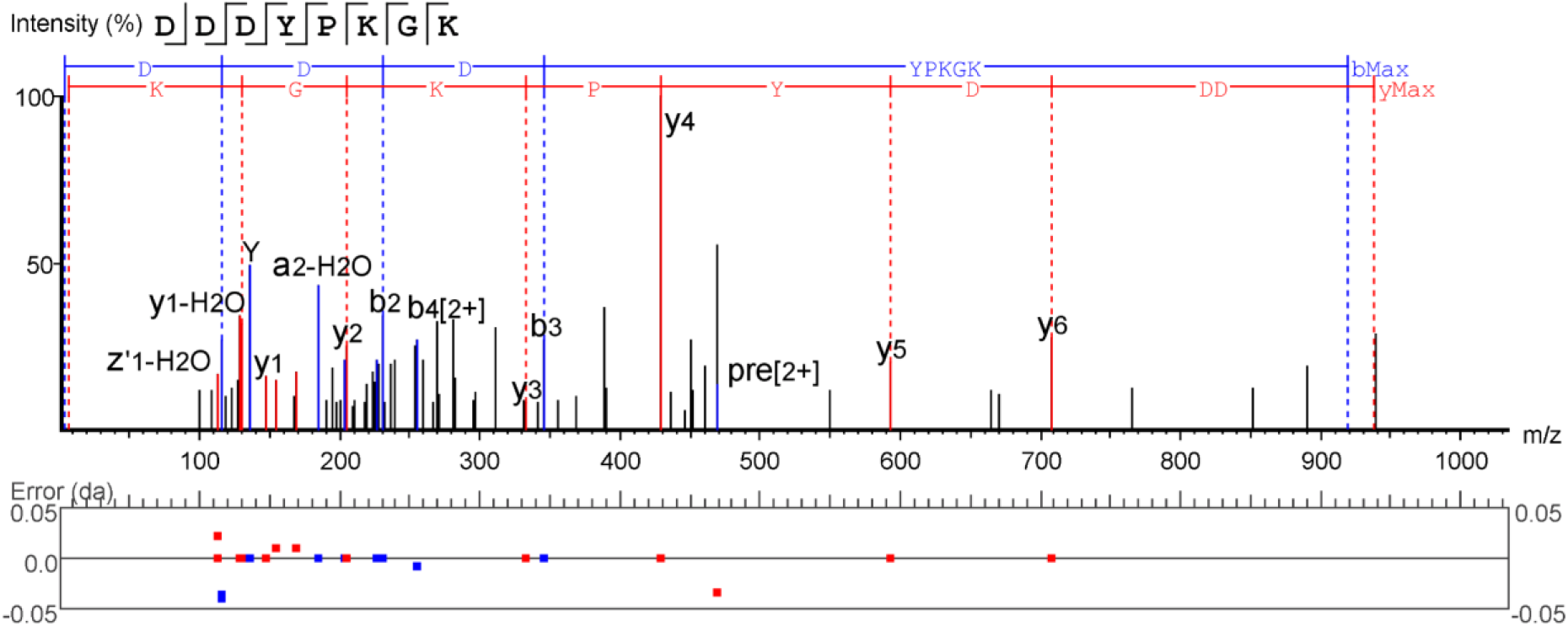
Annotated product ion spectrum, *m/z* = 469.2212

**Figure S10.**
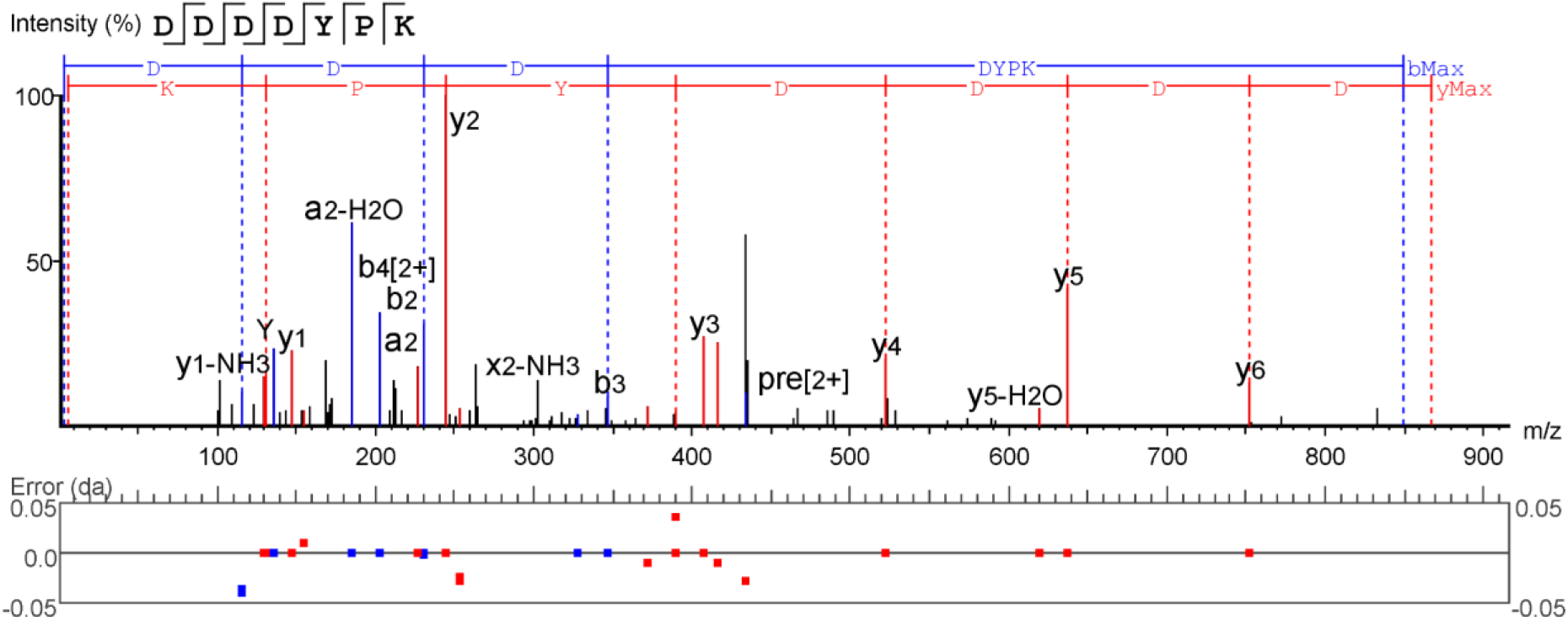
Annotated product ion spectrum, *m/z* = 434.1741

**Figure S11.**
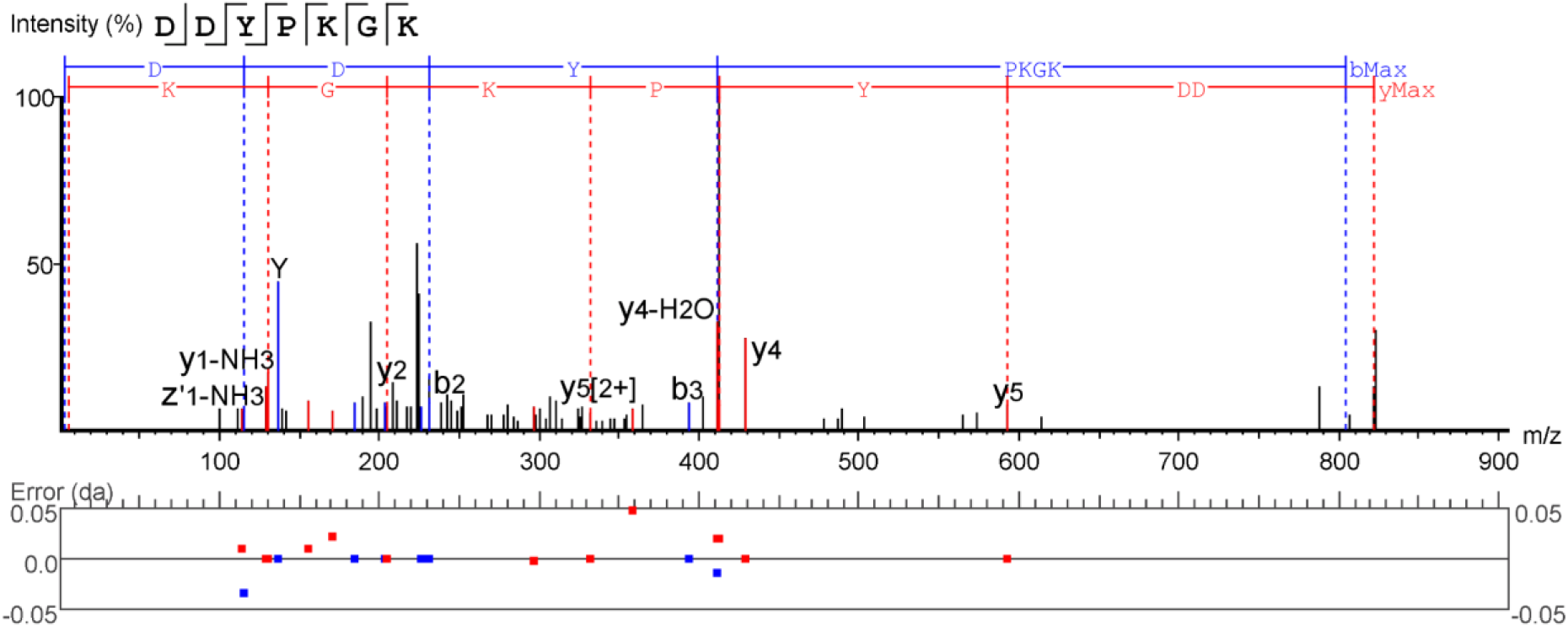
Annotated product ion spectrum, *m/z* = 411.7059

